# Single-cell nanodroplet processing proteomics pipeline for analysis of human-derived microglia

**DOI:** 10.1101/2025.10.02.680067

**Authors:** Ashley N. Ives, James M. Fulcher, Alex R. Bautista, Reta Birhanu Kitata, Sarah M. Williams, David A. Bennett, Philip L. De Jager, Vladislav A. Petyuk

## Abstract

Single-cell omics tools provide unique insights into heterogeneous cell populations and their responses to stimuli. For example, single-cell RNA sequencing has identified several transcriptionally distinct populations of microglia, which are resident immune cells of the central nervous system (CNS) that are responsive to CNS injury, infection, and neurodegeneration. To date, single cell studies of microglia have focused on RNA-sequencing or cytometry by time of flight (CyTOF) which provide indirect readouts of protein abundance or quantification of a limited number of targets. Herein, we present a workflow based on FACS-assisted isolation, cryopreservation and nanodroplet-based processing for single-cell mass spectrometry proteomics analysis of the postmortem human brain cortex-derived microglia. From a single microglial cell, 1039 proteins could be identified on average. As a proof-of-principle we applied single-cell proteomics for exploring the heterogeneity of brain microglia at the cellular level. This pilot proteomics data partially recapitulates the prior microglia subtypes. Specifically, we determined mitochondrial proteins, in particular members of NADH dehydrogenase (Complex I), cytochrome b-c1 (Complex III), cytochrome c oxidase (Complex IV), F1-ATPase (Complex V), and Na+/K+-ATPase complex drive variation across microglia. This pipeline offers the potential for identifying functionally and analytically relevant protein targets for microglia in Alzheimer’s disease and other neurological disorders.

**Significance of Study:** Microglia are a key brain cell type that may contribute to pathogenesis in neurodegenerative disease. Transcriptomic profiling of microglia from the central nervous system of humans and animal models has identified several subtypes of microglia, and complementary proteomic profiling of microglia is likely to provide functionally and therapeutically relevant targets, however single-proteomics studies of human-derived microglia are lacking. This work describes a label-free, single-cell proteomics approach for microglia isolated by fluorescence-activated cell sorting from a human donor that yields comparable numbers of identifications in comparison to prior single-cell RNA sequencing studies of microglia. This approach holds promise for enabling large-scale proteomics-based subtyping of microglia and studying their roles in neurodegenerative diseases.

## Introduction

Microglia are resident immune cells of the central nervous system (CNS) that dynamically respond to CNS injury, infection, and tissue maintenance^[1,2]^. The advent of single-cell omics techniques has revealed that microglia are highly diverse, heterogenous and dynamic, with many transcriptionally subtypes observed in both mouse and human subjects^[3–14]^. Subtypes of microglia that are associated with neurodegeneration and Alzheimer’s disease (AD) have recently been identified in single-cell RNA-Seq studies^[4,13,15,16]^, and it is anticipated that mapping these subtypes will better inform future diagnoses and treatment^[17,18]^. While these transcriptomic studies have provided important insights, prior bulk tissue omics studies demonstrate that proteomic and transcriptomic assays identify different disease-associated features in AD studies^[19]^. Additionally many neuropathologies are intrinsically protein-driven, including the formation of Aβ plaques, neurofibrillary tangles, Lewy bodies/ Lewy neurites, and TDP-43 inclusions^[20,21]^. Therefore, single-cell proteomics (scProteomics) may provide valuable insights when characterizing microglial heterogeneity.

To date, scProteomic studies of microglia have been limited and utilized targeted techniques (e.g. multiplexed ion beam imaging or cytometry based on time of flight) that focus on select proteins^[7,22–27]^, therefore, a method to more comprehensively characterize single microglial proteomes is highly desired^[28–31]^, and would provide new insights into the pathology of AD and other neurological disorders. Herein, we demonstrate a proof-of-principle and a feasibility of a label-free scProteomics workflow to study protein heterogeneity of the microglia population at the single-cell level isolated from a human brain cortex^[32,33]^. The workflow includes fluorescence-activated cell sorting (FACS)-assisted isolation of the microglia from freshly refrigerated tissue, cryopreservation, piezoelectric cell sorting (via cellenONE), nanodroplet processing platform (nanoPOTS)^[34]^, and high field asymmetric waveform ion mobility spectrometry (FAIMS)-data-independent acquisition (DIA) on the Orbitrap Astral mass analyzer^[35–38]^. Applying this approach to 48 single microglia from isolated from a post-mortem human brain cortex, we could identify 1039 ± 283 (mean ± s.d.) proteins per single cell before imputation. Our results show that the proteomics data captures the variation within microglia that in-principle agrees with prior results obtained using RNA-Seq in which various modes of metabolism were a primary axis separating microglia subtypes^[16]^; herein, mitochondrial proteins were confirmed as one of the key factors explaining the heterogeneity of the microglia cells. Given the encouraging results from this pilot study, we anticipate scProteomics will become a valuable tool for further refinement of the microglia subtypes, development of the surface protein markers and drug targets that can manage microglia subpopulations.

## Experimental Procedures

### Materials

Deionized water (18.2MΩ) was purified using a Barnstead Nanopure Infinity system (Los Angeles, CA, USA). N-dodecyl-β-D-maltoside (DDM), iodoacetamide (IAA), ammonium bicarbonate (ABC), and formic acid (FA) were obtained from Sigma (St. Louis, MO, USA). Trypsin (Promega, Madison, WI, USA) and Lys-C (Wako, Japan) were dissolved in 50 mM ABC before usage.

Dithiothreitol (DTT, No-Weigh format), acetonitrile (ACN) with 0.1% FA, water with 0.1% FA (MS grade), Hibernate-A, B27 supplement, GlutaMAX supplement, HBSS, RPMI-1640, EDTA, penicillin-streptomycin, and fetal bovine serum were purchased from Thermo Fisher Scientific (Waltham, MA, USA). CD11b MicroBeads, MACS LS columns, and QuadroMACS separator were purchased from Miltenyi Biotec (Bergisch Gladbach, Germany). Percoll was purchased from Cytiva (Marlborough, MA, USA). Cell staining buffer, anti-CD11b Alex Fluor 488 antibodies, and anti-CD45 Alexa Fluor 647 antibodies were purchased from BioLegend (San Diego, CA, USA). 7-AAD antibodies were purchased from BD Biosciences (Franklin Lakes, NJ, USA). HMC3 cells and EMEM were purchased from ATCC (Manassas, VA, USA).

### Fabrication of NanoPOTs Chips

The nanoPOTs chips were fabricated using photolithography, wet etching, and silanization as described previously^[34]^. The nanoSPLITs format was used^[39]^, with each chip containing 48 (4 x12) nanowells with a well diameter of 1.2 mm and an inter-well distance of 4.5 mm. Chip fabrication utilized a 25 mm x 75 mm glass slide pre-coated with chromium and photoresist (Telic Company, Valencia, USA). After photoresist exposure, development, and chromium etching (Transene Company, Inc., Danvers, USA), the exposed region was etched to a depth of ∼5 μm with buffered hydrofluoric acid. The freshly etched slide was dried by heating it at 120 °C for 1 h and then treated with oxygen plasma for 3 min (AP-300, Nordson March, Concord, USA). Heptadecafluoro-1,1,2,2-tetrahydrodecyl-dimethylchlorosilane (PFDS, Gelest, Germany) in 2,2,4-trimethylpentane (2:98 v/v) was applied onto the chip surface and incubated for 30 min to allow for silanization. The remaining chromium covering the wells was removed with chromium etchant, leaving elevated hydrophilic nanowells surrounded by a hydrophobic surface. Polyetheretherketone (PEEK) chip covers were machined as chip covers to provide sealing and protection of the droplets from evaporation during sample processing. Chips were wrapped in aluminum foil for long-term storage and intermediate steps during sample preparation.

### Source of Human Brain Sample

The autopsy brain specimen originated from brain donation programs at Rush University Medical Center/Rush Alzheimer’s Disease Center (Dr. David A. Bennet) in Chicago, USA. All procedures and research protocols were approved by the corresponding ethical committees of our collaborator’s institutions as well as the Institutional Review Board (IRB) of Columbia University Medical Center (protocol AAAR4962). The sampled brain specimen was taken from the medial frontal cortex region from an 83-year-old, white (non-hispanic), female donor. Autopsy was performed NS/SD with a PMI of 7.22 hours to fortified media. After weighing, the tissue was placed in ice-cold transportation medium (Hibernate-A medium containing 1% B27 serum-free supplement and 1% GlutaMax) and shipped overnight at 4 °C with priority shipping (media components from Thermo Fisher Scientific, Waltham, MA, USA).

### Microglia Isolation via Fluorescence-Activated Cell Sorting

The isolation of microglia was performed as previously published^[4,5]^. All stages of the protocol were carried out on ice. Briefly, tissue samples were placed in HBSS (Thermo Fisher Scientific, Waltham, MA, USA), weighed, and dissociated using a 15 mL glass Dounce homogenizer (Duran Wheaton Kimble) in increments of 0.5 g. The homogenate was filtered through a 100 μm cell strainer and then spun down at 450 RCF for 10 min. The pellet was resuspended in DPBS and myelin debris was removed via Percoll (Cytiva, Marlborough, MA, USA) gradient centrifugation at 3000 RCF for 10 min. The cell suspension was washed of Percoll using DPBS and subsequently centrifuged at 400 RCF for 10 min.

The resulting cell pellet was counted using a cell counter (Nexcelom) and then enriched for CD11b+ expression via magnetic-activated cell sorting (MACS) using anti-CD11b microbeads (Miltenyi, Gladbach, Germany), according to the manufacturer’s instructions. The cells were incubated with anti-CD11b magnetic beads for 15 min at 4 °C and subsequently washed in MACS buffer (PBS, 0.5% BSA, 2mM EDTA). The CD11b+ cells were purified on a LS MACS column (Miltenyi, Gladbach, Germany) attached to a QuadroMACS separator (Miltenyi, Gladbach, Germany). After washing the columns three times with MACS buffer, the CD11b+ cells were flushed from the column with MACS buffer and pelleted.

The CD11b+ enriched cell pellet was then incubated with anti-CD11b Alexa Fluor 488 (BioLegend, San Diego, CA, USA), anti-CD45 Alexa Fluor 647 (BioLegend, San Diego, CA, USA), and 7-AAD (BD Biosciences, Franklin Lakes, New Jersey) antibodies for 20 min on ice. Subsequently the cell suspension was washed twice with staining buffer (BioLegend, San Diego, CA, USA), filtered through a 70 μm filter, and the CD11b+/CD45+/7AAD− cells were sorted via fluorescence-activated cell sorting (FACS) using a BD Influx cell sorter. All sorting was performed using a 100-µm nozzle. The sorting speed was kept between 8,000 and 10,000 events per second. Following FACS, the cell suspensions were exchanged into RPMI-1640 (Thermo Fisher Scientific, Waltham, MA, USA) supplemented with 40% FBS (Thermo Fisher Scientific, Waltham, MA, USA) and 10% DMSO (Bio-Techne, Minneapolis, MN, USA) and stored at –80 °C. The samples were shipped for analysis on dry ice and again stored at –80 °C.

### Preparation of HMC3 Samples

Human microglial clone 3 (CRL-3304, ATCC, Manassas, VA, USA) cells were cultured in Eagle’s Minimum Essential Medium (ATCC, Manassas, VA, USA), supplemented with 10% FBS and 1% penicillin-streptomycin (Thermo Fisher Scientific, Waltham, MA, USA), and incubated in a humidified 5% CO_2_ incubator at 37 °C. 0.75-1×10^6^ HMC3 cells were seeded into a 10 cm dish (Corning, Corning, NY, USA) and passaged every second day with new culture medium. The cells were collected for analysis at early (4-7) passages. Cell suspensions were exchanged into RPMI-1640 supplemented with 40% FBS and 10% DMSO and shipped on dry ice. Samples were stored at –80 °C.

### Sorting and Deposition of the Individual Cells onto Nanodroplet Processing Chips

Cell suspensions were centrifuged at 500 RCF for 5 min and the supernatant was removed and replaced with 1xPBS. This step was repeated three times. A CellenONE instrument equipped with a glass piezo capillary (P-20-CM) was utilized for single cell isolation. Sorting parameters for microglia included a pulse length of 51 μs, a nozzle voltage of 76 V, a frequency of 500 Hz, an LED delay of 200 μs, and an LED pulse of 3 μs. Sorting parameters for HMC3 included a pulse length of 49 μs, a nozzle voltage of 74 V, a frequency of 500 Hz, an LED delay of 200 μs, and an LED pulse of 3 μs. The slide stage was operated at dew point control mode to reduce droplet evaporation and keep protoplasts at low temperature after sorting (2-4 °C). Single cells were isolated based on diameter and elongation to exclude cellular debris and non-dissociated cell clumps. Specific criteria for microglia selection included cell diameters of 17 to 36 μm, maximum circularity of 2, and maximum elongation of 2.18. Specific criteria for HMC3 selection included cell diameters of 26 to 39 μm, maximum circularity of 2, and maximum elongation of 2.19. All single cells were sorted based on bright field images in real time. After sorting, all chips were wrapped in parafilm and aluminum foil before being frozen and stored at –80 °C. Additionally, all single cell sorting events were manually inspected using the brightfield images to remove any non-intact cells or potential doublets from downstream analysis.

### NanoPOTS-based Sample Processing

All liquid dispensing was performed using the CellenONE instrument equipped with a glass piezo capillary (PDC 60, No Coating). For scProteomics with nanoPOTS chips, extraction was accomplished by dispensing 150 nL of extraction buffer containing 50 mM ABC, 0.1% DDM, diluted PBS (x0.5), and 2 mM DTT and incubating the chip at 60 °C for 60 min. Denatured and reduced proteins were alkylated through the addition of 50 nL 15 mM IAA before incubation for 30 min in darkness at room temperature. Alkylated proteins were then digested by adding 50 nL 50 mM ABC with 0.01 ng/nL of Lys-C and 0.04 ng/nL of trypsin and incubating at 37 °C overnight. The digestion reaction was then quenched by adding 50 nL of 5% formic acid before drying the chip under vacuum at room temperature. All chips were wrapped in parafilm and aluminum foil and stored at –80 °C until LC-MS/MS analysis.

### LC-MS/MS Instrumental Analysis

Samples prepared on nanoPOTS chips were transferred to 384-well plates by adding 5 uL 0.1% formic acid per nanoPOTs well, overlaying the chips and well plates, and centrifuging at 3000 RCF for 1 min. This process was repeated three times. Samples were analyzed using a Neo UHPLC system (Thermo Fisher Scientific, Waltham, MA, USA) coupled to an Orbitrap Astral mass spectrometer (Thermo Fisher Scientific, Waltham, MA, USA) equipped with a FAIMS Pro interface (Thermo Fisher Scientific, Waltham, MA, USA) and an EASY-Spray source. Source settings included a spray voltage of 2.2 kV, ion transfer tube temperature of 280 °C, carrier gas flow of 3.5 L/min, and compensation voltage of –40 V. Liquid chromatography was performed using a Vanquish Neo LC (Thermo Fisher Scientific, Waltham, MA, USA), running a 70 SPD (samples per day) separation method, with each separation having a 14 min active gradient and 6 min for sample loading and column equilibration. The Vanquish Neo was configured to run in the trap-and-elute mode, utilizing the PepMap Neo Trap Cartridge (Thermo Fisher Scientific, Waltham, MA, USA) for sample trapping and reverse flow onto the analytical column. A PepMap ES906 analytical column was used for reverse phase elution of the peptides and heated to 45 °C during operation. 5 uL of sample were used for each injection, with mobile phases consisting of buffer A (0.1% formic acid in water) and buffer B (0.1% formic acid in acetonitrile). The exact gradient details and flow rates are described in **Table S1**. The Orbitrap Astral was operated in data-independent acquisition mode (DIA). MS1 spectra were recorded using a resolution of 120K, scan range of 400-900 m/z, 1 microscan, automated gate control (AGC) target of 5e6, and a maximum injection time of 356 ms. For MS2, nonoverlapping isolation windows 20 m/z in width were scanned across 400-900 m/z. MS2 spectra were recorded using 1 microscan, an ACG target of 8e5, maximum injection time of 40 ms, and a HCD collision energy of 25%.

### Database Search Parameters and Acceptance Criteria for Identifications

All proteomic data raw files were processed using Spectronaut (version 20.1.250624.92449) and searched against the *Homo sapiens* UniProt protein sequence database (UP000005640; includes 20,664 genes accessed on 02/25). The common Repository of FBS Proteins (cRFP) was also added to the search database and includes 210 bovine proteins^[40]^, along with enzyme contaminants such as porcine trypsin and endoprotease Lys-C from *Lysobacter enzymogenes*. For DirectDIA+ (library-free) searches replicates were defined as the same condition in the search settings. Allowable peptide lengths were restricted to 7-52 amino acids with 2 allowable missed cleavages. Carbamidomethylation of cysteines was listed a static modification.

Methionine oxidation and N-terminal protein acetylation were listed as variable modifications. Factory settings were used for all analyses and for library generation including a precursor q-value cutoff of 0.01, precursor posterior error probability (PEP) cutoff of 0.2, protein q-value cutoff (experiment) of 0.01, protein q-value cutoff (run) of 0.05, and protein PEP cutoff of 0.75. PSM, peptide, and protein group FDRs were kept at 0.01.

Precursors were exported from Spectronaut and filtered for a q-value of <0.01. Precursors that could not be unambiguously assigned as human were removed to prevent incorrect quantitation of human proteins due to possible bovine serum contamination. Precursors corresponding to keratins were removed as contaminants. Single cell datasets with <500 identified precursors were removed from further analysis. Precursors with < 50% sample coverage were removed prior to imputation^[41]^. The protein log2-intensities were converted to a relative scale by subtracting the median intensity for each protein. Imputation was performed using svdImpute method from the “pcaMethods” Bioconductor R package^[42]^. Precursor’s log2-transformed abundances were then summarized at the protein level using the “fast_MaxLFQ” function from the “iq” R package^[43,44]^. Data was then batch corrected for individual nanoPOTs chips using the ComBat method from the “sva” package^[45]^. The median relative abundance of the histone proteins was then calculated for all samples and then applied as a normalization factor as previously described^[46,47]^. Gene ontology (GO) biological process terms were assigned to identified proteins using the “gprofiler2” R package^[48]^. To identify the biological pathways explaining the diversity of the microglial cells, we summarized the protein data at the GO term level using “rrollup” function in the “MSnSet.utils” R package and followed by principal component analysis. Intensity Based Absolute Quantification (iBAQ) values were calculated by summing all peptide intensities for a given protein and dividing this value by the number of theoretically observable tryptic peptides between 6 and 50 amino acids in length^[49]^.

## Results

### Coverage of the Microglia Proteome by scProteomics

The scProteomics platform utilized piezoelectric sorting of cell suspensions, trypsin digestion on the nanoPOTs chips, followed by LC-MS/MS on the Orbitrap Astral mass spectrometer with the FAIMS Pro interface (**Figure 1A**). Without imputation, we obtained 3290 ± 1059 (mean ± s.d.) precursors (defined as modified peptide and charge state) and 1039 ± 283 proteins from single microglia (N = 47) using a library-free search (**Figures 1B-1C**). We obtained higher identifications from single cells of a human, microglia-like immortalized cell line (Human Microglial Cell 3, HMC3) with 7483 ± 2427 precursors and 2009 ± 538 proteins (N = 46 single cells). Precursors were relatively complete across single cell measurements with 2922 precursors (50%) observed in at least half of single microglia measurements (**Figure S1**). The precursor completeness was higher in single HMC3 with 72% of precursors being observed in at least 50% of single cell measurements (**Figure S2**). We also observed several microglia-specific markers including CD45 (Gene symbol: PTPRC), CD11b (Gene symbol: ITGAM), P2RY12, and HEXB (data not shown)^[50]^. Additionally, 5% of the observed proteins (190 proteins) from single cell microglia have previously been reported to be enriched in aged microglia (the HuMi_Aged gene set)^[5]^; the intensity Based Absolute Quantification (iBAQ) values for the top 500 most abundant proteins are shown as **Figure 2**.

**Figure 1.**
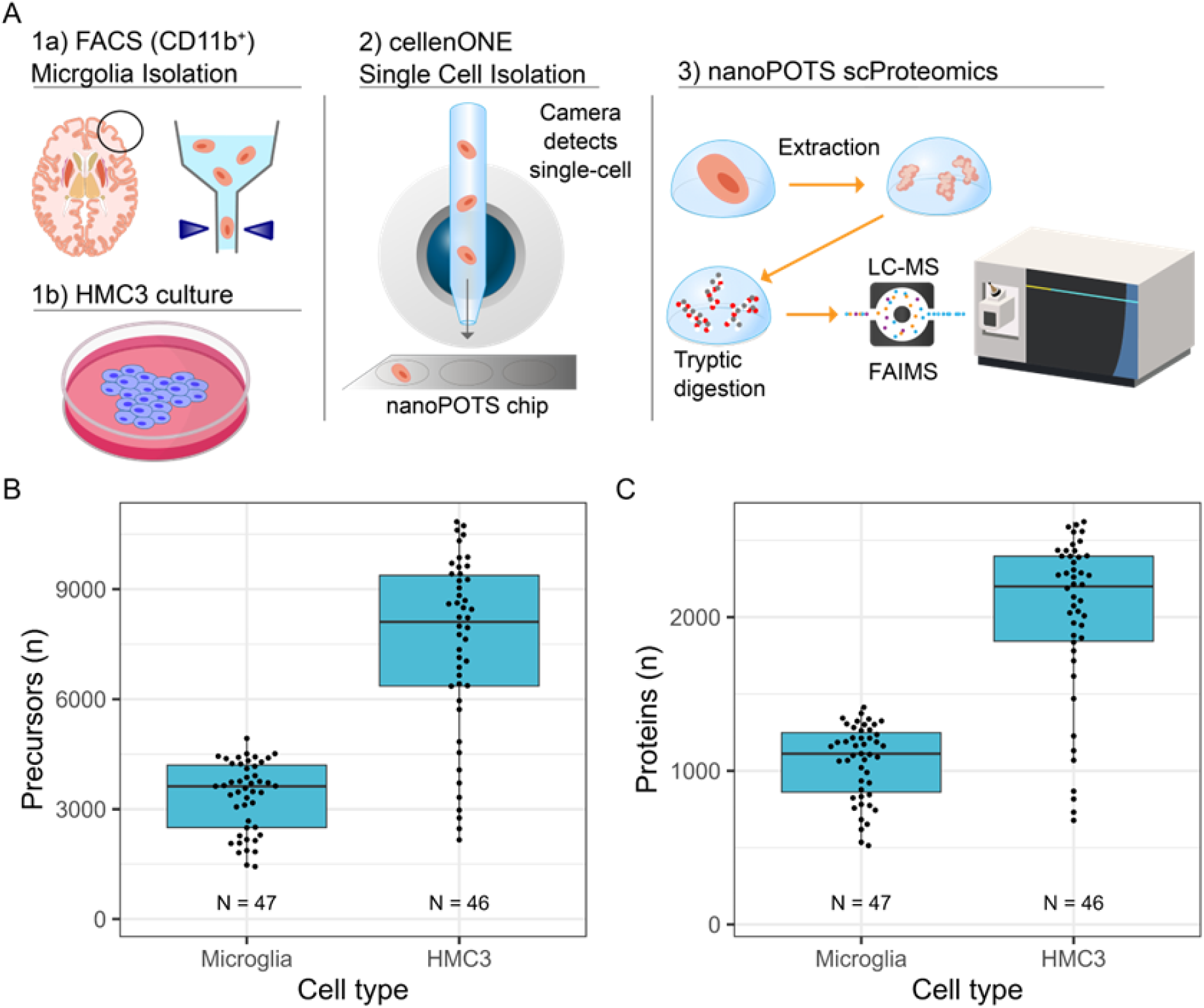
(A) scProteomics workflow. Cell suspensions are processed into single cell samples using piezoelectric cell sorting, digested using nanoPOTS sample preparation, analyzed using FAIMS-based MS data-independent acquisition method, and searched using Spectronaut 20. (B) Precursor– and (C) protein-level identifications before imputation for microglia and HMC3. The ‘N’ indicates the number of measured single cells.

**Figure 2.**
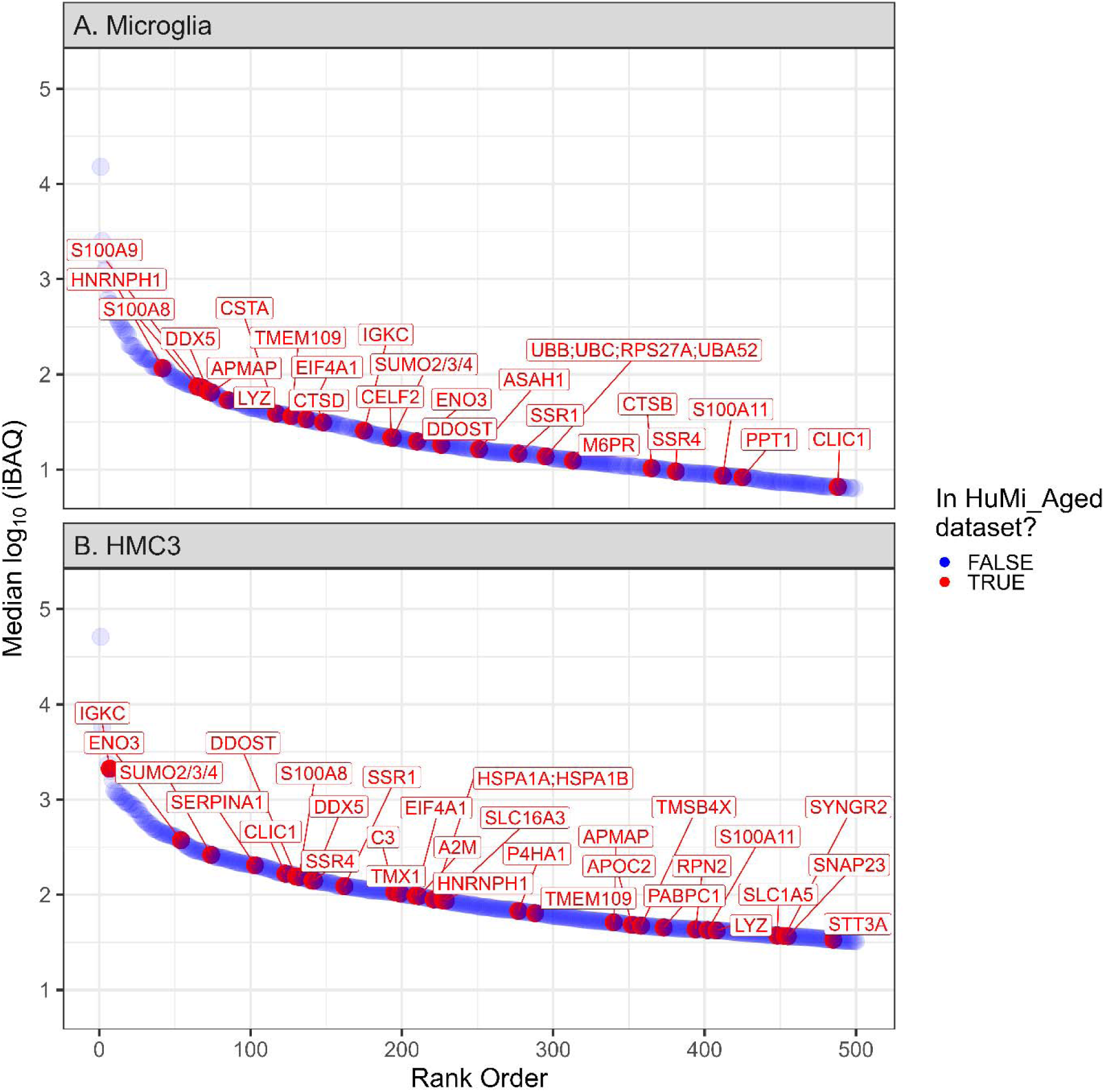
Identified proteins from single microglia and HMC3. Median log10(intensity Based Absolute Quantification) for proteins observed in single (A) microglia and (B) HMC3. Intensities are before imputation or normalization and have been filtered for the top 500 most abundant proteins that are observed in >50% of single cell samples. Proteins are plotted in rank order from most to least abundant. Proteins listed in the HuMi_Aged dataset are denoted in red with gene symbols annotated, while all other proteins are denoted in blue.

### Proteome Heterogeneity of Microglia and HMC3 Cells

Following batch correction (**Figures S3-S4**) and imputation, we applied principal component analysis (PCA) to explore proteins and gene ontology (GO) terms that explain the most variance in the data. For microglia, the top contributing proteins to PC1 include TUBB2A, TUBA4A, CAMK2A, RBFOX3, and ANP32E while proteins that contribute to PC2 include ATP15F1D, SLC25A4, PPT1, COX6C, ATP1A2, and NDUFA4 (**Figures 3A-3B**). From this principal component analysis, we observed that single cells seemed to group based on cell size (diameter). We further explored how cell size correlates with top principal components (PCs1-10) by calculating Pearson correlation coefficients using the eigencor function from PCAtools. This analysis revealed that PC1 is significantly associated with cell diameter for single cell microglia (**Figure S5**). Aggregating proteins to GO biological process terms followed by a similar PCA analysis revealed mitochondrial-related processes including respiration, mitochondrial ATP synthesis, and oxidative phosphorylation as the top pathways varying throughout the microglia population (**Figures 3C-3D**), consistent with prior RNA-Seq profiles^[16]^. Proteins aggregated into these mitochondrial terms include protomers of NADH dehydrogenase (Complex I), cytochrome b-c1 (Complex III), cytochrome c oxidase (Complex IV), F1-ATPase (Complex V), and Na+/K+-ATPase complex (**Supplementary File 2**). Other terms with high contribution to principal components include microtubule-related processes and mitosis, which are largely driven by several tubulins. This principal component analysis was repeated for HMC3; top contributing proteins including various histones, SNRPB/N, SRSF7, APOC1, and IPO4 (**Figure 4A-4B**). Aggregation to GO biological processes reveals several terms driven by eukaryotic initiation factors (e.g. formation of cytoplasmic translation initiation complex, viral translational termination-reinitiation), apolipoproteins (high-density lipoprotein particle remodeling), and various complement factors or immunoglobulin chains (complement activation) (**Figure 4C-4D, Supplementary File 3**). Finally, we found that cell diameter was significantly associated with only the third principal component in HMC3 cells (**Figure S5**).

**Figure 3.**
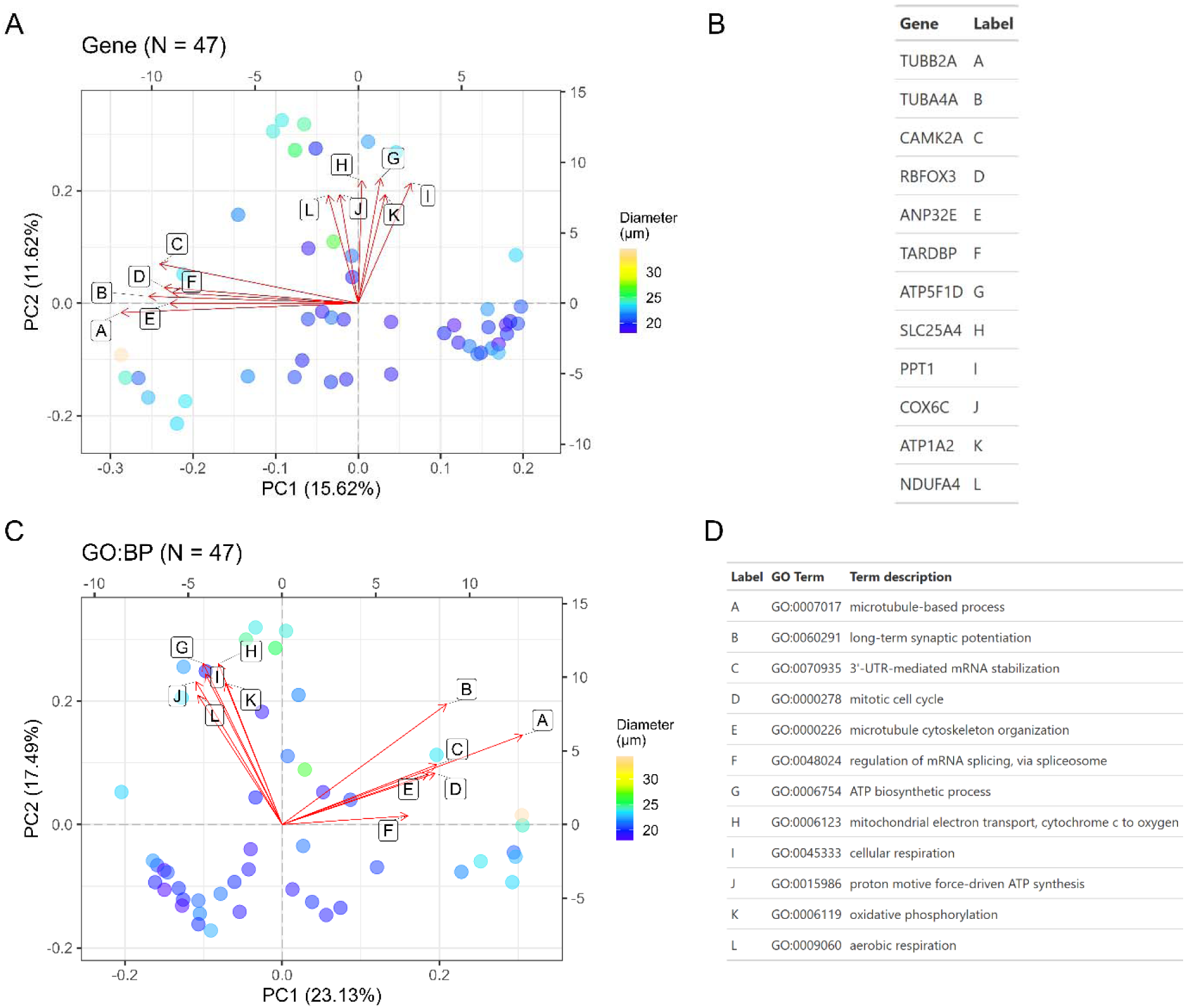
Principal component analysis (PCA) of single cell microglia. Each dot represents a single cell measurement with color fill denoting the cell diameter (µm) as determined by the CellenONE camera. (A) PCA based on label-free protein quantities (DirectDIA+ analysis). Loadings are represented by vectors and indicate top genes that contribute to cell variability in a specific direction. (B) Gene identities for biplot loadings shown in panel (A). (C) PCA analysis following aggregation of label-free protein quantities to gene ontology (GO) terms, specifically for biological processes. Loadings are represented by vectors and indicate top GO terms that contribute to cell variability in a specific direction. Explained variance per PC is listed in the axis labels. (D) GO biological processes for biplot loadings shown in panel (C).

**Figure 4.**
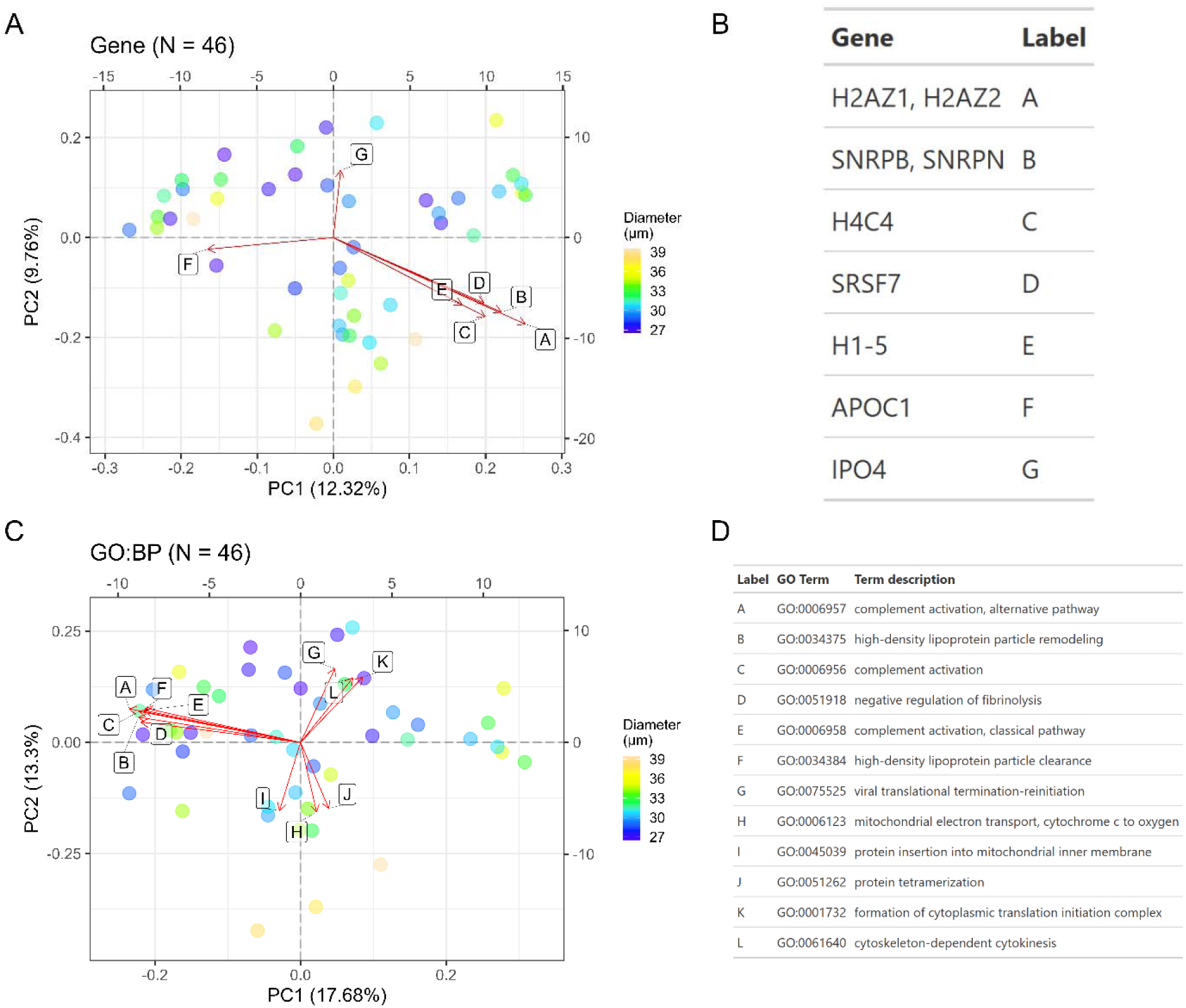
Principal component analysis (PCA) of single cell HMC3. Each dot represents a single cell measurement with color fill denoting the cell diameter (µm) as determined by the CellenONE camera. (A) PCA based on label-free protein quantities (DirectDIA+ analysis). Loadings are represented by vectors and indicate top genes that contribute to cell variability in a specific direction. (B) Gene identities for biplot loadings shown in panel (A). (C) PCA analysis following aggregation of label-free protein quantities to gene ontology (GO) terms, specifically for biological processes. Loadings are represented by vectors and indicate top GO terms that contribute to cell variability in a specific direction. Explained variance per PC is listed in the axis labels. (D) GO biological processes for biplot loadings shown in panel (C).

## Discussion

### Proteome Coverage

Our work represents the first application of label-free scProteomics to the individual microglia cells isolated from post-mortem human brain cortex. Microglia are challenging given that viable cells rapidly decline after death, with noticeable changes in gene expression occurring within 6 hours postmortem delay^[51,52]^. Given the small cell size and thus limited protein mass presents another challenge. *Ex vivo* microglia are estimated to have about 10-fold less protein than postmortem-cultured microglia or immortalized cell lines^[53]^. Using FACS enrichment followed by piezoelectric cellenONE sorting and nanoPOTS techniques, we demonstrated results comparable to single-cell RNA sequencing based on number of genes identified per single microglia; in prior studies the median number of identified genes is <1500 per donor^[4]^. This nanoPOTs approach is also compatible with same-cell measurements of transcriptomes and proteomes (nanoSPLITs)^[39]^ and spatial scProteomics coupled with laser capture microdissection. Thus, it opens avenues for future exploration of transcriptional and spatial heterogeneity in the context of microglia subtyping.

### Proteome Variability in Single Cell Microglia Population

Our results show that mitochondrial proteins partially explain the variation between individual microglia. Microglia undergo changes in energy metabolism during activation and differentiation^[54]^. Resting microglia primarily use oxidative phosphorylation, while pro-inflammatory populations rely on aerobic glycolysis and anti-inflammatory populations depend on mitochondrial oxidation^[55–57]^. Our results could be capturing these metabolic shifts given that GO terms contributing to variation include genes involved in respiration, proton transmembrane transport, and mitochondrial ATP synthesis. Additionally, we found that cytoskeletal proteins (e.g. tubulins) also contributed to variation and roughly aligned with separation of cells by diameter. This agrees with prior transcriptomic observations that cytoskeletal proteins correlate with cell size in mammalian cell lines^[58]^. For microglia, we found that the first principal component for protein-level dimension reduction significantly correlated with cell diameter highlighting the importance of considering cell size as a phenotypic variable in future analyses. This is consistent with prior observations that cell size is one of the major contributors to variation in single-cell proteomes^[46]^. Cell size contributed less to the variation in HMC3, only associating with the third principal component. Additionally, the total variance explained by first and second principal components in HMC3 is lower than in microglia, suggesting that the immortalized microglia (HMC3) are more homogeneous in their proteomic landscape.

## Conclusion

We describe a pipeline for scProteomic analysis of single microglia isolated from the human brain. The steps included enrichment of microglia using CD11 marker, further FACS-assisted enrichment, cryopreservation, re-sorting and deposition onto nanoPOTs chips, followed by scProteomics processing pipeline. Quantification of the abundances of ∼1000 proteins per cell on average was achieved, placing the approach at the same level as single-cell RNA-seq. This orthogonal and direct approach for measuring global protein abundances in single-cells will complement existing transcriptomic studies by revealing distinct protein-expression patterns that are crucial for understanding microglia phenotypes and subtypes, especially in the context of neurodegenerative diseases.

## Supporting information

Supp_1.docx

## Acknowledgements

This work was supported by the NIH National Institute awards U01 AG061356 (P.L.D.J. and D.A.B.), R01 AG015819 (D.A.B.), R01 AG17917 (D.A.B.), P30 AG10161 (D.A.B.), and P30 AG72975 (D.A.B.). We thank Dr. Matthew Monroe for his assistance in depositing the raw proteomic data onto MassIVE. We thank Dr. David Ross Hall for guidance on using Spectronaut.

## Conflict of Interest Statement

The authors have declared no conflict of interest.

## Data Availability

The mass spectrometry raw data have been deposited to the ProteomeXchange Consortium via the MassIVE partner repository with dataset identifier MSV000099294.

## Authors’ Contributions

V.A.P. and P.L.D.J. conceptualized and designed the initial project. D.A.B. supplied the donor tissue. A. R. B. performed the HMC3 cell culturing and FACS-sorting. A.N.I. and J.M.F. performed the LC-MS/MS sample preparation. J.M.F and S.M.W. maintained and operated instrumentation for LC-MS analysis. A.N.I, J.M.F., and V.A.P. analyzed data. A.N.I. and V.A.P. drafted the manuscript. All other authors reviewed, provided constructive feedback, and approved the final manuscript.

## Abbreviations

ABC: Ammonium Bicarbonate
AGC: Automatic Gain Control
DDM: n-Dodecyl-β-D665 Maltoside
DIA: Data-Independent Acquisition
DTT: Dithiothreitol
FA: Formic Acid
FACS: Fluorescence-Activated Cell Sorting
FAIMS: High Field Asymmetric Waveform Ion Mobility Spectrometry
FDR: False Discovery Rate
HCD: Higher-Energy Collisional Dissociation
IAA: Iodoacetamide
LFQ: Label Free Quantification
nanoPOTS: Nanodroplet Processing in One Pot for Trace Samples
PBS: Phosphate-Buffered Saline
PEEK: Polyetheretherketone
PFDS: Heptadecafluoro-1,1,2,2-Tetrahydrodecyl-Dimethylchlorosilane
scProteomics: Single-Cell Proteomics

## Notes

### Competing Interest Statement

The authors have declared no competing interest.

## References

[1] Colonna, M., & Butovsky, O. (2017). Microglia Function in the Central Nervous System During Health and Neurodegeneration. Annual Review of Immunology, 35(Volume 35, 2017), 441–468. 10.1146/annurev-immunol-051116-052358

[2] Nayak, D., Roth, T. L., & McGavern, D. B. (2014). Microglia Development and Function*. Annual Review of Immunology, 32(Volume 32, 2014), 367–402. 10.1146/annurev-immunol-032713-120240

[3] Masuda, T., Sankowski, R., Staszewski, O., & Prinz, M. (2020). Microglia Heterogeneity in the Single-Cell Era. Cell Reports, 30(5), 1271–1281. 10.1016/j.celrep.2020.01.010

[4] Olah, M., Menon, V., Habib, N., Taga, M. F., Ma, Y., Yung, C. J., Cimpean, M., Khairallah, A., Coronas-Samano, G., Sankowski, R., Grün, D., Kroshilina, A. A., Dionne, D., Sarkis, R. A., Cosgrove, G. R., Helgager, J., Golden, J. A., Pennell, P. B., Prinz, M., … De Jager, P. L. (2020). Single cell RNA sequencing of human microglia uncovers a subset associated with Alzheimer’s disease. Nature Communications, 11(1), 6129. 10.1038/s41467-020-19737-2

[5] Olah, M., Patrick, E., Villani, A.-C., Xu, J., White, C. C., Ryan, K. J., Piehowski, P., Kapasi, A., Nejad, P., Cimpean, M., Connor, S., Yung, C. J., Frangieh, M., McHenry, A., Elyaman, W., Petyuk, V., Schneider, J. A., Bennett, D. A., De Jager, P. L., & Bradshaw, E. M. (2018). A transcriptomic atlas of aged human microglia. Nature Communications, 9(1), 539. 10.1038/s41467-018-02926-5

[6] Li, Q., Cheng, Z., Zhou, L., Darmanis, S., Neff, N. F., Okamoto, J., Gulati, G., Bennett, M. L., Sun, L. O., Clarke, L. E., Marschallinger, J., Yu, G., Quake, S. R., Wyss-Coray, T., & Barres, B. A. (2019). Developmental Heterogeneity of Microglia and Brain Myeloid Cells Revealed by Deep Single-Cell RNA Sequencing. Neuron, 101(2), 207–223.e10. 10.1016/j.neuron.2018.12.006

[7] Sankowski, R., Böttcher, C., Masuda, T., Geirsdottir, L., Sagar, null, Sindram, E., Seredenina, T., Muhs, A., Scheiwe, C., Shah, M. J., Heiland, D. H., Schnell, O., Grün, D., Priller, J., & Prinz, M. (2019). Mapping microglia states in the human brain through the integration of high-dimensional techniques. Nature Neuroscience, 22(12), 2098– 2110. 10.1038/s41593-019-0532-y

[8] Hammond, T. R., Dufort, C., Dissing-Olesen, L., Giera, S., Young, A., Wysoker, A., Walker, A. J., Gergits, F., Segel, M., Nemesh, J., Marsh, S. E., Saunders, A., Macosko, E., Ginhoux, F., Chen, J., Franklin, R. J. M., Piao, X., McCarroll, S. A., & Stevens, B. (2019). Single-Cell RNA Sequencing of Microglia throughout the Mouse Lifespan and in the Injured Brain Reveals Complex Cell-State Changes. Immunity, 50(1), 253–271.e6. 10.1016/j.immuni.2018.11.004

[9] Masuda, T., Sankowski, R., Staszewski, O., Böttcher, C., Amann, L., Sagar, Scheiwe, C., Nessler, S., Kunz, P., van Loo, G., Coenen, V. A., Reinacher, P. C., Michel, A., Sure, U., Gold, R., Grün, D., Priller, J., Stadelmann, C., & Prinz, M. (2019). Spatial and temporal heterogeneity of mouse and human microglia at single-cell resolution. Nature, 566(7744), 388–392. 10.1038/s41586-019-0924-x

[10] Yaqubi, M., Groh, A. M. R., Dorion, M.-F., Afanasiev, E., Luo, J. X. X., Hashemi, H., Sinha, S., Kieran, N. W., Blain, M., Cui, Q.-L., Biernaskie, J., Srour, M., Dudley, R., Hall, J. A., Sonnen, J. A., Arbour, N., Prat, A., Stratton, J. A., Antel, J., & Healy, L. M. (2023). Analysis of the microglia transcriptome across the human lifespan using single cell RNA sequencing. Journal of Neuroinflammation, 20(1), 132. 10.1186/s12974-023-02809-7

[11] Sousa, C., Golebiewska, A., Poovathingal, S. K., Kaoma, T., Pires-Afonso, Y., Martina, S., Coowar, D., Azuaje, F., Skupin, A., Balling, R., Biber, K., Niclou, S. P., & Michelucci, A. (2018). Single-cell transcriptomics reveals distinct inflammation-induced microglia signatures. EMBO Reports, 19(11), e46171. 10.15252/embr.201846171

[12] Mathys, H., Adaikkan, C., Gao, F., Young, J. Z., Manet, E., Hemberg, M., De Jager, P. L., Ransohoff, R. M., Regev, A., & Tsai, L.-H. (2017). Temporal Tracking of Microglia Activation in Neurodegeneration at Single-Cell Resolution. Cell Reports, 21(2), 366–380. 10.1016/j.celrep.2017.09.039

[13] Keren-Shaul, H., Spinrad, A., Weiner, A., Matcovitch-Natan, O., Dvir-Szternfeld, R., Ulland, T. K., David, E., Baruch, K., Lara-Astaiso, D., Toth, B., Itzkovitz, S., Colonna, M., Schwartz, M., & Amit, I. (2017). A Unique Microglia Type Associated with Restricting Development of Alzheimer’s Disease. Cell, 169(7), 1276–1290.e17. 10.1016/j.cell.2017.05.018

[14] Sunna, S., Bowen, C. A., Ramelow, C. C., Santiago, J. V., Kumar, P., & Rangaraju, S. (2023). Advances in proteomic phenotyping of microglia in neurodegeneration. PROTEOMICS, 23(13–14), 2200183. 10.1002/pmic.202200183

[15] Green, G. S., Fujita, M., Yang, H.-S., Taga, M., Cain, A., McCabe, C., Comandante-Lou, N., White, C. C., Schmidtner, A. K., Zeng, L., Sigalov, A., Wang, Y., Regev, A., Klein, H.-U., Menon, V., Bennett, D. A., Habib, N., & De Jager, P. L. (2024). Cellular communities reveal trajectories of brain ageing and Alzheimer’s disease. Nature, 633(8030), 634– 645. 10.1038/s41586-024-07871-6

[16] Tuddenham, J. F., Taga, M., Haage, V., Marshe, V. S., Roostaei, T., White, C., Lee, A. J., Fujita, M., Khairallah, A., Zhang, Y., Green, G., Hyman, B., Frosch, M., Hopp, S., Beach, T. G., Serrano, G. E., Corboy, J., Habib, N., Klein, H.-U., … De Jager, P. L. (2024). A cross-disease resource of living human microglia identifies disease-enriched subsets and tool compounds recapitulating microglial states. Nature Neuroscience, 27(12), 2521–2537. 10.1038/s41593-024-01764-7

[17] Miao, J., Ma, H., Yang, Y., Liao, Y., Lin, C., Zheng, J., Yu, M., & Lan, J. (2023). Microglia in Alzheimer’s disease: pathogenesis, mechanisms, and therapeutic potentials. Frontiers in Aging Neuroscience, 15. 10.3389/fnagi.2023.1201982

[18] Hansen, D. V., Hanson, J. E., & Sheng, M. (2018). Microglia in Alzheimer’s disease. The Journal of Cell Biology, 217(2), 459–472. 10.1083/jcb.201709069

[19] Johnson, E. C. B., Carter, E. K., Dammer, E. B., Duong, D. M., Gerasimov, E. S., Liu, Y., Liu, J., Betarbet, R., Ping, L., Yin, L., Serrano, G. E., Beach, T. G., Peng, J., De Jager, P. L., Haroutunian, V., Zhang, B., Gaiteri, C., Bennett, D. A., Gearing, M., … Seyfried, N. T. (2022). Large-scale deep multi-layer analysis of Alzheimer’s disease brain reveals strong proteomic disease-related changes not observed at the RNA level. Nature Neuroscience, 25(2), 213–225. 10.1038/s41593-021-00999-y

[20] Jo, M., Lee, S., Jeon, Y.-M., Kim, S., Kwon, Y., & Kim, H.-J. (2020). The role of TDP-43 propagation in neurodegenerative diseases: integrating insights from clinical and experimental studies. Experimental & Molecular Medicine, 52(10), 1652–1662. 10.1038/s12276-020-00513-7

[21] Zhang, J., Zhang, Y., Wang, J., Xia, Y., Zhang, J., & Chen, L. (2024). Recent advances in Alzheimer’s disease: mechanisms, clinical trials and new drug development strategies. Signal Transduction and Targeted Therapy, 9(1), 1–35. 10.1038/s41392-024-01911-3

[22] Mrdjen, D., Amouzgar, M., Cannon, B., Liu, C., Spence, A., McCaffrey, E., Bharadwaj, A., Tebaykin, D., Bukhari, S., Hartmann, F. J., Kagel, A., Vijayaragavan, K., Oliveria, J. P., Yakabi, K., Serrano, G. E., Corrada, M. M., Kawas, C. H., Camacho, C., Bosse, M., … Bendall, S. C. (2023). Spatial proteomics reveals human microglial states shaped by anatomy and neuropathology. Research Square, rs.3.rs-2987263. 10.21203/rs.3.rs-2987263/v1

[23] Böttcher, C., Schlickeiser, S., Sneeboer, M. A. M., Kunkel, D., Knop, A., Paza, E., Fidzinski, P., Kraus, L., Snijders, G. J. L., Kahn, R. S., Schulz, A. R., Mei, H. E., Hol, E. M., Siegmund, B., Glauben, R., Spruth, E. J., de Witte, L. D., & Priller, J. (2019). Human microglia regional heterogeneity and phenotypes determined by multiplexed single-cell mass cytometry. Nature Neuroscience, 22(1), 78–90. 10.1038/s41593-018-0290-2

[24] Ajami, B., Samusik, N., Wieghofer, P., Ho, P. P., Crotti, A., Bjornson, Z., Prinz, M., Fantl, W. J., Nolan, G. P., & Steinman, L. (2018). Single-cell mass cytometry reveals distinct populations of brain myeloid cells in mouse neuroinflammation and neurodegeneration models. Nature Neuroscience, 21(4), 541–551. 10.1038/s41593-018-0100-x

[25] Mrdjen, D., Pavlovic, A., Hartmann, F. J., Schreiner, B., Utz, S. G., Leung, B. P., Lelios, I., Heppner, F. L., Kipnis, J., Merkler, D., Greter, M., & Becher, B. (2018). High-Dimensional Single-Cell Mapping of Central Nervous System Immune Cells Reveals Distinct Myeloid Subsets in Health, Aging, and Disease. Immunity, 48(2), 380–395.e6. 10.1016/j.immuni.2018.01.011

[26] Vijayaragavan, K., Cannon, B. J., Tebaykin, D., Bossé, M., Baranski, A., Oliveria, J. P., Bukhari, S. A., Mrdjen, D., Corces, M. R., McCaffrey, E. F., Greenwald, N. F., Sigal, Y., Marquez, D., Khair, Z., Bruce, T., Goldston, M., Bharadwaj, A., Montine, K. S., Angelo, R. M., … Bendall, S. C. (2022). Single-cell spatial proteomic imaging for human neuropathology. Acta Neuropathologica Communications, 10(1), 158. 10.1186/s40478-022-01465-x

[27] Miedema, A., Wijering, M. H. C., Eggen, B. J. L., & Kooistra, S. M. (2020). High-Resolution Transcriptomic and Proteomic Profiling of Heterogeneity of Brain-Derived Microglia in Multiple Sclerosis. Frontiers in Molecular Neuroscience, 13. 10.3389/fnmol.2020.583811

[28] Sunna, S., Bowen, C. A., Ramelow, C. C., Santiago, J. V., Kumar, P., & Rangaraju, S. (2023). Advances in proteomic phenotyping of microglia in neurodegeneration. Proteomics, 23(13–14), e2200183. 10.1002/pmic.202200183

[29] Davis, E., & Lloyd, A. F. (2024). The proteomic landscape of microglia in health and disease. Frontiers in Cellular Neuroscience, 18. 10.3389/fncel.2024.1379717

[30] Goto-Silva, L., & Junqueira, M. (2021). Single-cell proteomics: A treasure trove in neurobiology. Biochimica et Biophysica Acta (BBA)-Proteins and Proteomics, 1869(7), 140658. 10.1016/j.bbapap.2021.140658

[31] Sankowski, R., Monaco, G., & Prinz, M. (2022). Evaluating microglial phenotypes using single-cell technologies. Trends in Neurosciences, 45(2), 133–144. 10.1016/j.tins.2021.11.001

[32] Janabi, N., Peudenier, S., Héron, B., Ng, K. H., & Tardieu, M. (1995). Establishment of human microglial cell lines after transfection of primary cultures of embryonic microglial cells with the SV40 large T antigen. Neuroscience Letters, 195(2), 105–108. 10.1016/0304-3940(94)11792-H

[33] Dello Russo, C., Cappoli, N., Coletta, I., Mezzogori, D., Paciello, F., Pozzoli, G., Navarra, P., & Battaglia, A. (2018). The human microglial HMC3 cell line: where do we stand? A systematic literature review. Journal of Neuroinflammation, 15(1), 259. 10.1186/s12974-018-1288-0

[34] Zhu, Y., Piehowski, P. D., Zhao, R., Chen, J., Shen, Y., Moore, R. J., Shukla, A. K., Petyuk, V. A., Campbell-Thompson, M., Mathews, C. E., Smith, R. D., Qian, W.-J., & Kelly, R. T. (2018). Nanodroplet processing platform for deep and quantitative proteome profiling of 10-100 mammalian cells. Nature Communications, 9(1), 882. 10.1038/s41467-018-03367-w

[35] Bubis, J. A., Arrey, T. N., Damoc, E., Delanghe, B., Slovakova, J., Sommer, T. M., Kagawa, H., Pichler, P., Rivron, N., Mechtler, K., & Matzinger, M. (2025). Challenging the Astral mass analyzer to quantify up to 5,300 proteins per single cell at unseen accuracy to uncover cellular heterogeneity. Nature Methods, 22(3), 510–519. 10.1038/s41592-024-02559-1

[36] Sweet, S., Chain, D., Yu, W., Martin, P., Rebelatto, M., Chambers, A., Cecchi, F., & Kim, Y. J. (2022). The addition of FAIMS increases targeted proteomics sensitivity from FFPE tumor biopsies. Scientific Reports, 12(1), 13876. 10.1038/s41598-022-16358-1

[37] Swearingen, K. E., & Moritz, R. L. (2012). High-field asymmetric waveform ion mobility spectrometry for mass spectrometry-based proteomics. Expert Review of Proteomics, 9(5), 505–517. 10.1586/epr.12.50

[38] Stejskal, K., Op de Beeck, J., Dürnberger, G., Jacobs, P., & Mechtler, K. (2021). Ultrasensitive NanoLC-MS of Subnanogram Protein Samples Using Second Generation Micropillar Array LC Technology with Orbitrap Exploris 480 and FAIMS PRO. Analytical Chemistry, 93(25), 8704–8710. 10.1021/acs.analchem.1c00990

[39] Fulcher, J. M., Markillie, L. M., Mitchell, H. D., Williams, S. M., Engbrecht, K. M., Degnan, D. J., Bramer, L. M., Moore, R. J., Chrisler, W. B., Cantlon-Bruce, J., Bagnoli, J. W., Qian, W.-J., Seth, A., Paša-Tolić, L., & Zhu, Y. (2024). Parallel measurement of transcriptomes and proteomes from same single cells using nanodroplet splitting. Nature Communications, 15(1), 10614. 10.1038/s41467-024-54099-z

[40] Shin, J., Kwon, Y., Lee, S., Na, S., Hong, E. Y., Ju, S., Jung, H.-G., Kaushal, P., Shin, S., Back, J. H., Choi, S. Y., Kim, E. H., Lee, S. J., Park, Y. E., Ahn, H.-S., Ahn, Y., Kabir, M. H., Park, S.-J., Yang, W. S., … Lee, C. (2019). Common Repository of FBS Proteins (cRFP) To Be Added to a Search Database for Mass Spectrometric Analysis of Cell Secretome. Journal of Proteome Research, 18(10), 3800–3806. 10.1021/acs.jproteome.9b00475

[41] Webb-Robertson, B.-J. M., Wiberg, H. K., Matzke, M. M., Brown, J. N., Wang, J., McDermott, J. E., Smith, R. D., Rodland, K. D., Metz, T. O., Pounds, J. G., & Waters, K. M. (2015). Review, Evaluation, and Discussion of the Challenges of Missing Value Imputation for Mass Spectrometry-Based Label-Free Global Proteomics. Journal of Proteome Research, 14(5), 1993–2001. 10.1021/pr501138h

[42] Stacklies, W., Redestig, H., Scholz, M., Walther, D., & Selbig, J. (2007). pcaMethods—a bioconductor package providing PCA methods for incomplete data. Bioinformatics, 23(9), 1164–1167. 10.1093/bioinformatics/btm069

[43] Pham, T. V., Henneman, A. A., & Jimenez, C. R. (2020). iq: an R package to estimate relative protein abundances from ion quantification in DIA-MS-based proteomics. *Bioinformatics (Oxford*, England), 36(8), 2611–2613. 10.1093/bioinformatics/btz961

[44] Cox, J., Hein, M. Y., Luber, C. A., Paron, I., Nagaraj, N., & Mann, M. (2014). Accurate proteome-wide label-free quantification by delayed normalization and maximal peptide ratio extraction, termed MaxLFQ. Molecular & Cellular Proteomics: MCP, 13(9), 2513– 2526. 10.1074/mcp.M113.031591

[45] Johnson, W. E., Li, C., & Rabinovic, A. (2007). Adjusting batch effects in microarray expression data using empirical Bayes methods. Biostatistics, 8(1), 118–127. 10.1093/biostatistics/kxj037

[46] Lanz, M. C., Valenzuela, L. F., Elias, J. E., & Skotheim, J. M. (2023). Cell Size Contributes to Single-Cell Proteome Variation. Journal of Proteome Research, 22(12), 3773–3779. 10.1021/acs.jproteome.3c00441

[47] Lanz, M. C., Zhang, S., Swaffer, M. P., Ziv, I., Götz, L. H., Kim, J., McCarthy, F., Jarosz, D. F., Elias, J. E., & Skotheim, J. M. (2024). Genome dilution by cell growth drives starvation-like proteome remodeling in mammalian and yeast cells. Nature Structural & Molecular Biology, 31(12), 1859–1871. 10.1038/s41594-024-01353-z

[48] Kolberg, L., Raudvere, U., Kuzmin, I., Vilo, J., & Peterson, H. (2020). gprofiler2 –-an R package for gene list functional enrichment analysis and namespace conversion toolset g:Profiler. F1000Research, 9, ELIXIR-709. 10.12688/f1000research.24956.2

[49] Schwanhäusser, B., Busse, D., Li, N., Dittmar, G., Schuchhardt, J., Wolf, J., Chen, W., & Selbach, M. (2011). Global quantification of mammalian gene expression control. Nature, 473(7347), 337–342. 10.1038/nature10098

[50] Honarpisheh, P., Lee, J., Banerjee, A., Blasco-Conesa, M. P., Honarpisheh, P., d’Aigle, J., Mamun, A. A., Ritzel, R. M., Chauhan, A., Ganesh, B. P., & McCullough, L. D. (2020). Potential caveats of putative microglia-specific markers for assessment of age-related cerebrovascular neuroinflammation. Journal of Neuroinflammation, 17(1), 366. 10.1186/s12974-020-02019-5

[51] Heng, Y., Dubbelaar, M. L., Marie, S. K. N., Boddeke, E. W. G. M., & Eggen, B. J. L. (2021). The effects of postmortem delay on mouse and human microglia gene expression. Glia, 69(4), 1053–1060. 10.1002/glia.23948

[52] Mizee, M. R., Miedema, S. S. M., van der Poel, M., Adelia, Schuurman, K. G., van Strien, M. E., Melief, J., Smolders, J., Hendrickx, D. A., Heutinck, K. M., Hamann, J., & Huitinga, I. (2017). Isolation of primary microglia from the human post-mortem brain: effects of ante– and post-mortem variables. Acta Neuropathologica Communications, 5, 16. 10.1186/s40478-017-0418-8

[53] Lloyd, A. F., Martinez-Muriana, A., Davis, E., Daniels, M. J. D., Hou, P., Mancuso, R., Brenes, A. J., Sinclair, L. V., Geric, I., Snellinx, A., Craessaerts, K., Theys, T., Fiers, M., Strooper, B. D., & Howden, A. J. M. (2024). Deep proteomic analysis of microglia reveals fundamental biological differences between model systems. Cell Reports, 43(11). 10.1016/j.celrep.2024.114908

[54] Jurga, A. M., Paleczna, M., & Kuter, K. Z. (2020). Overview of General and Discriminating Markers of Differential Microglia Phenotypes. Frontiers in Cellular Neuroscience, 14, 198. 10.3389/fncel.2020.00198

[55] Orihuela, R., McPherson, C. A., & Harry, G. J. (2016). Microglial M1/M2 polarization and metabolic states. British Journal of Pharmacology, 173(4), 649–665. 10.1111/bph.13139

[56] Fumagalli, M., Lombardi, M., Gressens, P., & Verderio, C. (2018). How to reprogram microglia toward beneficial functions. Glia, 66(12), 2531–2549. 10.1002/glia.23484

[57] Moss, D. W., & Bates, T. E. (2001). Activation of murine microglial cell lines by lipopolysaccharide and interferon-gamma causes NO-mediated decreases in mitochondrial and cellular function. The European Journal of Neuroscience, 13(3), 529–538. 10.1046/j.1460-9568.2001.01418.x

[58] Miettinen, T. P., Pessa, H. K. J., Caldez, M. J., Fuhrer, T., Diril, M. K., Sauer, U., Kaldis, P., & Björklund, M. (2014). Identification of transcriptional and metabolic programs related to mammalian cell size. Current Biology: CB, 24(6), 598–608. 10.1016/j.cub.2014.01.071

