## Supplementary material for "Single-cell nanodroplet processing proteomics pipeline for analysis of human-derived microglia": Supp_1.docx

### Figure S1. Precursor completeness across single microglia.

Percentile bins containing precursors with different degrees of data completeness (number of cells observed in/ total single cell measurements) from single cell microglia (N = 47).

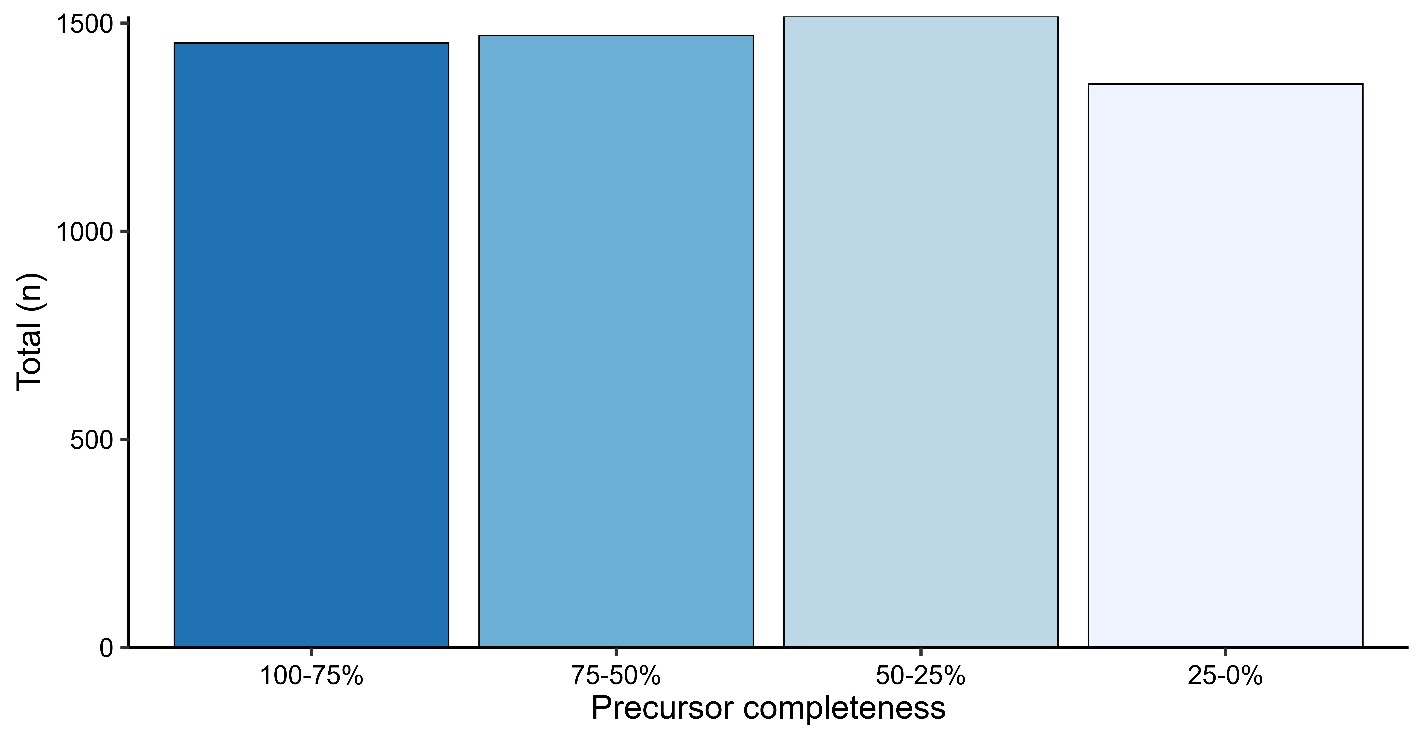

### Figure S2. Precursor completeness across single HMC3.

Percentile bins containing precursors with different degrees of data completeness (number of cells observed in/ total single cell measurements) from single cell HMC3 (N = 46).

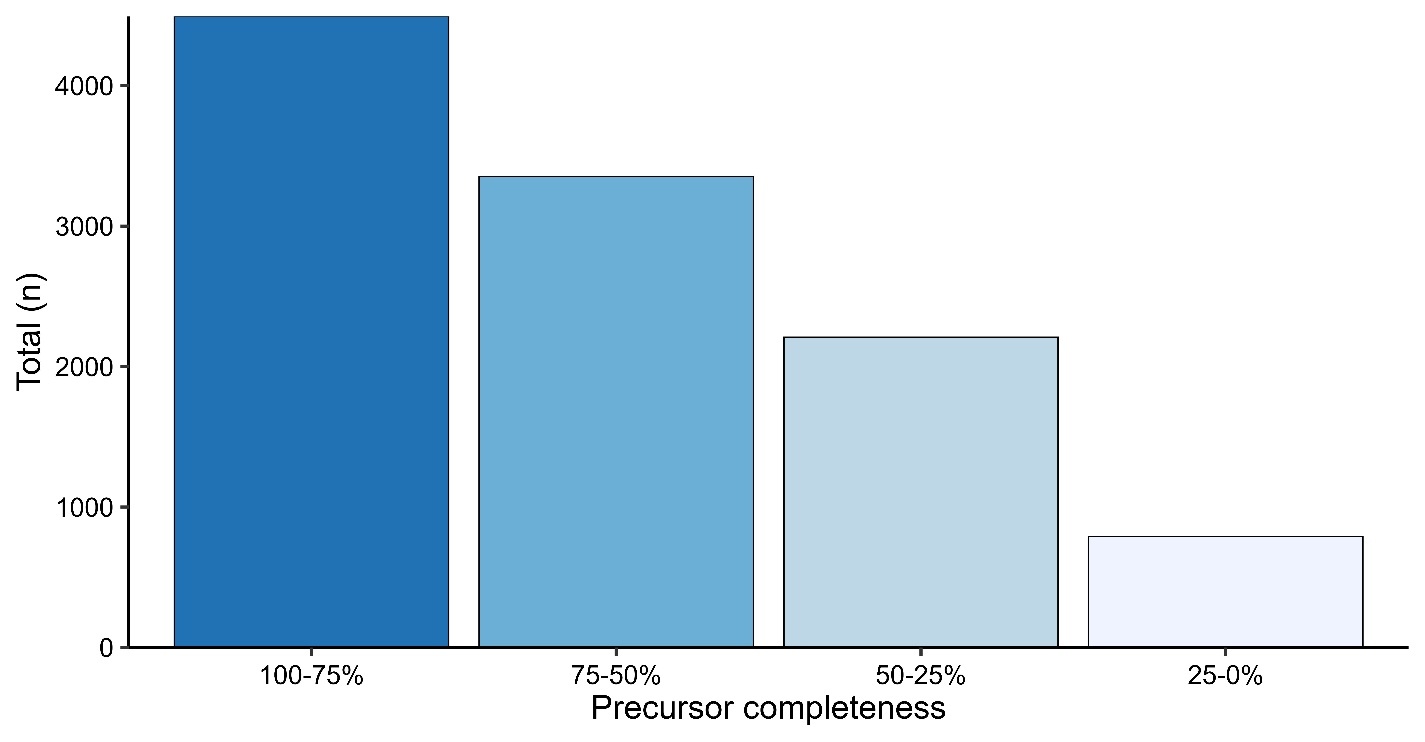

### Figure S3. Effect of batch correction on single microglia data.

PCA plots of maxLFQ protein abundances before A) and after B) batch correction using the ComBat procedure. Samples are colored according to the nanoPOTs chip (batch).

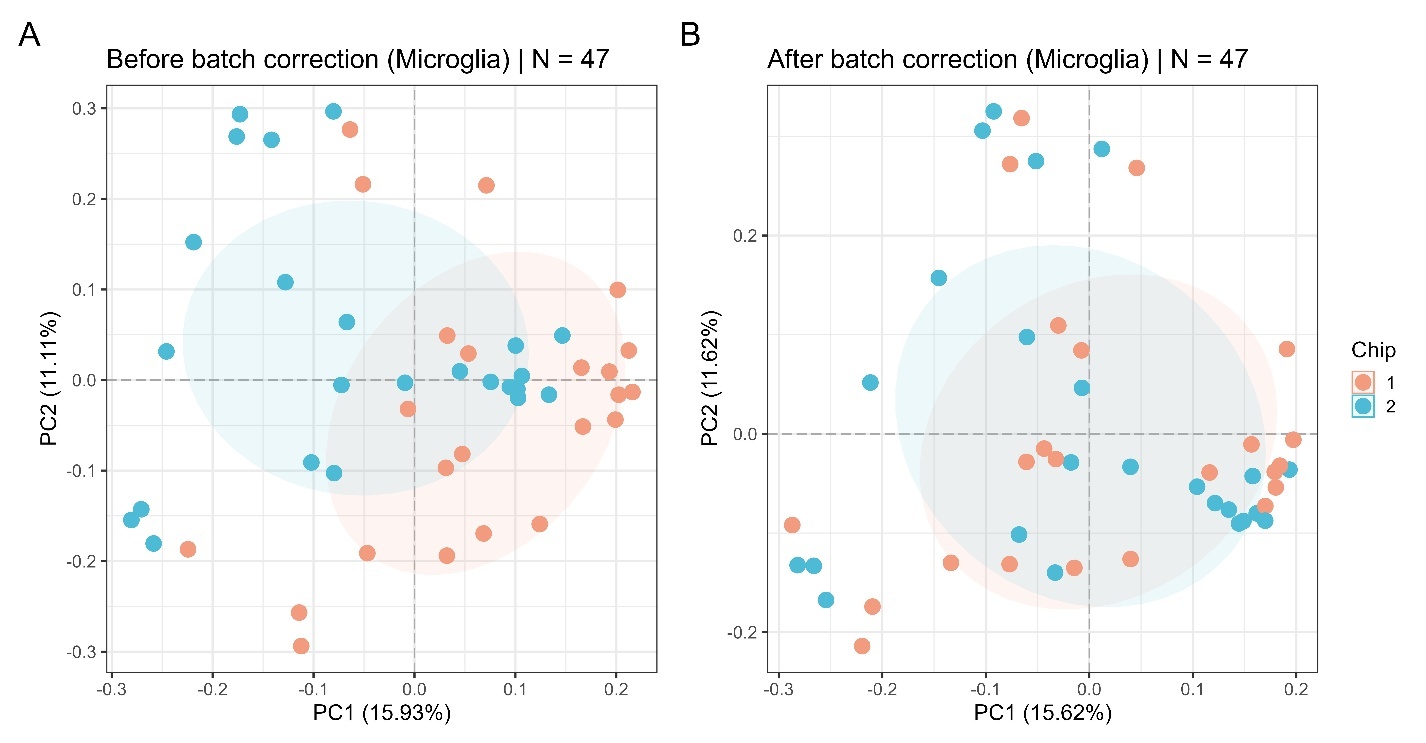

### Figure S4. Effect of batch correction on single HMC3 data.

PCA plots of maxLFQ protein abundances before A) and after B) batch correction using the ComBat procedure. Samples are colored according to the nanoPOTs chip (batch).

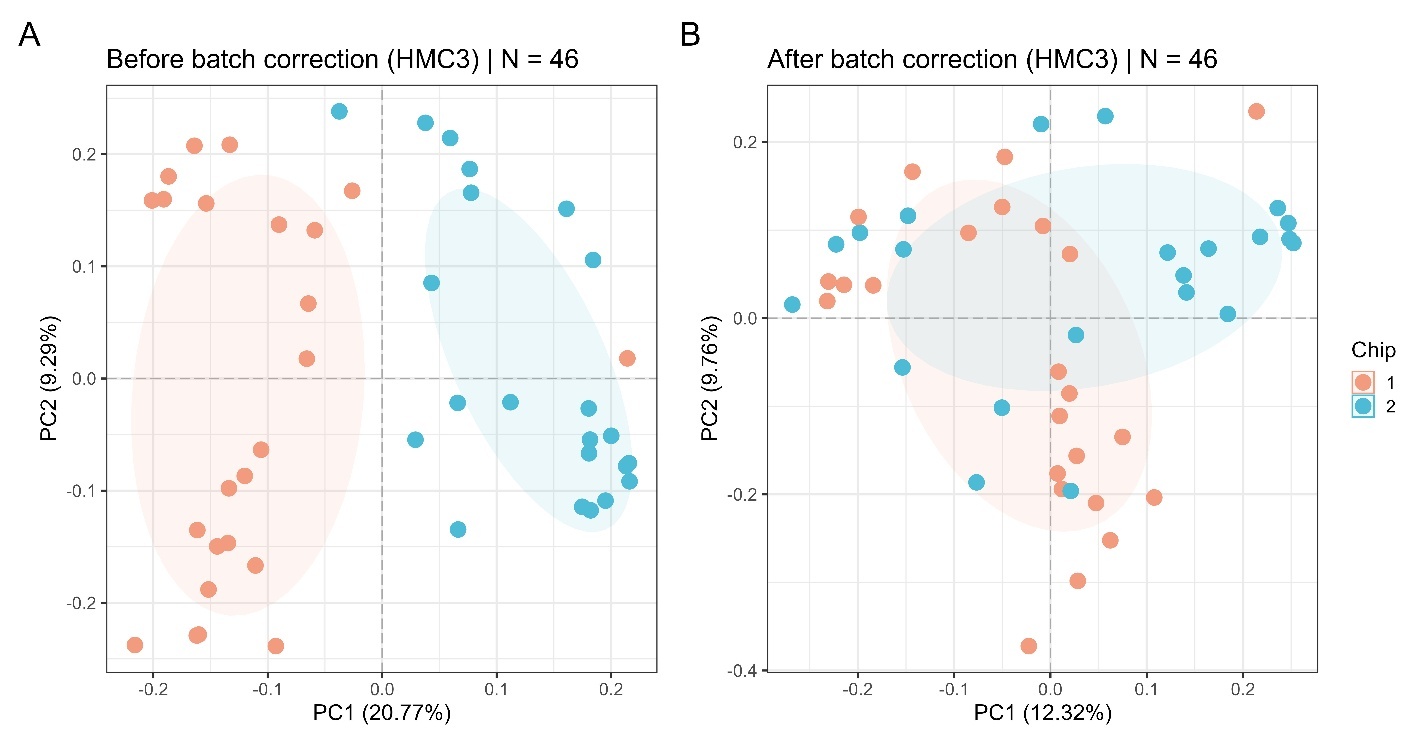

### Figure S5. Eigen correlation plots between principal components and cell diameter.

Pearson correlation coefficients were calculated between the top 10 principal components and cell diameter for single microglia and HMC3. Color fill denotes correlation magnitude and direction. Point size shows -log_10_(BH-adjusted p-values). Significant correlations are denoted with stars (p-values ≤0.01 = “**”).

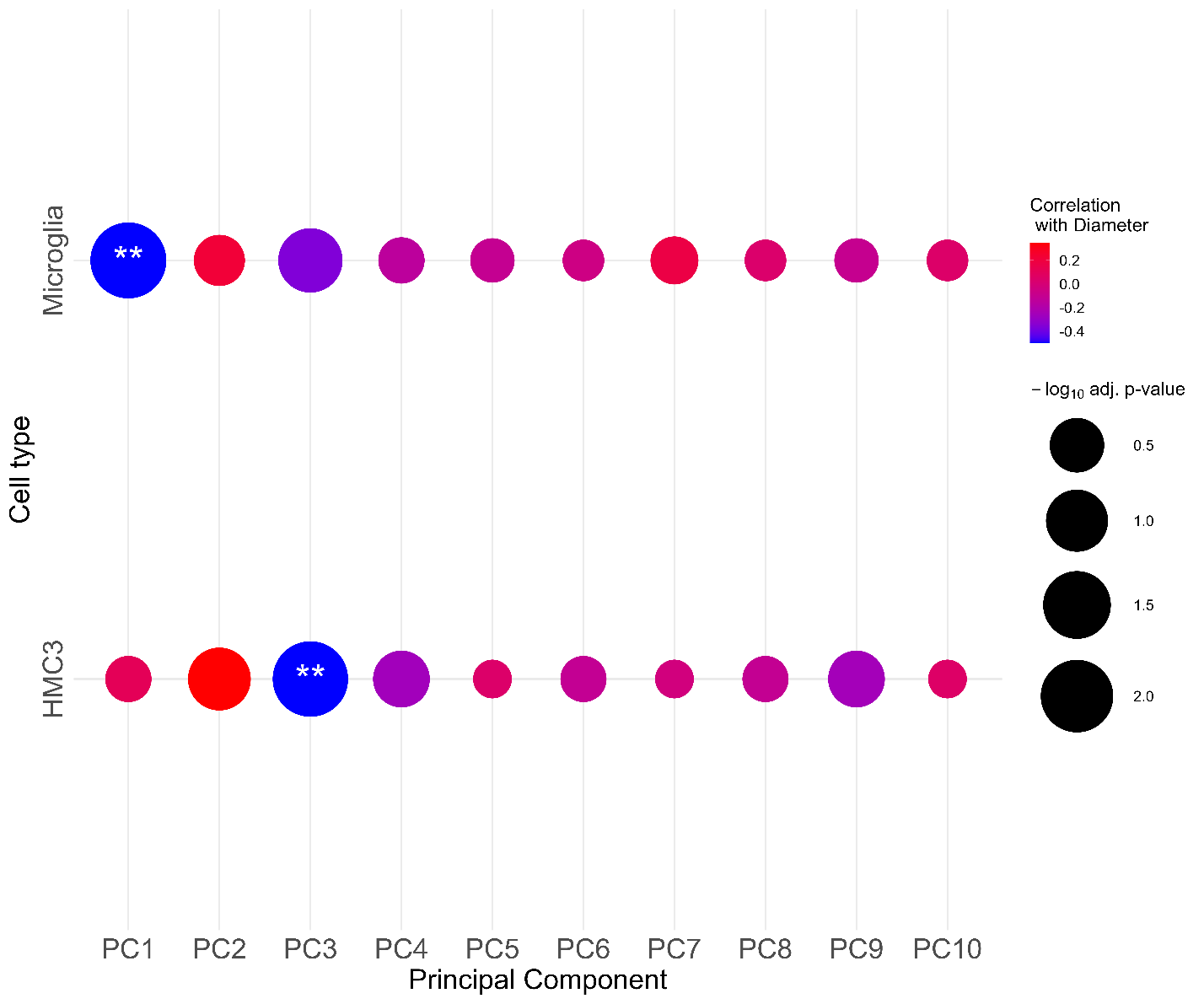

### Table S1. Liquid chromatography gradient for 70 SPD method.

| **Step** | **Time** | **Flow (uL/min)** | **%B** |
| --- | --- | --- | --- |
| Equilibration | 0.0 | 3.50 | 1.0 |
| Equilibration | 0.001 | 2.8 | 1.0 |
| Gradient | 0.501 | 1.30 | 4.0 |
| Gradient | 0.901 | 1.10 | 8.0 |
| Gradient | 12.301 | 1.10 | 22.5 |
| Gradient | 15.301 | 1.10 | 35.0 |
| Gradient | 15.501 | 1.10 | 55.0 |
| Wash | 16.001 | 2.8 | 99.0 |
| Wash | 17.201 | 2.8 | 99.0 |
| Wash | 18.000 | 2.8 | 1.0 |
