## Supplementary material for "Single-cell nanodroplet processing proteomics pipeline for analysis of human-derived microglia": Supp_2.pdf

| Accession | Gene | GO_ID | GO_Term |
|---|---|---|---|
| P07910 | HNRNPC | GO:0070935 | 3'-UTR-mediated mRNA stabilization |
| P98175 | RBM10 | GO:0070935 | 3'-UTR-mediated mRNA stabilization |
| Q13148 | TARDBP | GO:0070935 | 3'-UTR-mediated mRNA stabilization |
| Q13151 | HNRNPA0 | GO:0070935 | 3'-UTR-mediated mRNA stabilization |
| Q15717 | ELAVL1 | GO:0070935 | 3'-UTR-mediated mRNA stabilization |
| O75489 | NDUFS3 | GO:0009060 | aerobic respiration |
| O95168 | NDUFB4 | GO:0009060 | aerobic respiration |
| O96000 | NDUFB10 | GO:0009060 | aerobic respiration |
| P04040 | CAT | GO:0009060 | aerobic respiration |
| P14927 | UQCRB | GO:0009060 | aerobic respiration |
| P19404 | NDUFV2 | GO:0009060 | aerobic respiration |
| P28331 | NDUFS1 | GO:0009060 | aerobic respiration |
| P30049 | ATP5F1D | GO:0009060 | aerobic respiration |
| P31930 | UQCRC1 | GO:0009060 | aerobic respiration |
| P40926 | MDH2 | GO:0009060 | aerobic respiration |
| Q16718 | NDUFA5 | GO:0009060 | aerobic respiration |
| O75306 | NDUFS2 | GO:0009060 | aerobic respiration |
| P06576 | ATP5F1B | GO:0006754 | ATP biosynthetic process |
| P25705 | ATP5F1A | GO:0006754 | ATP biosynthetic process |
| P30049 | ATP5F1D | GO:0006754 | ATP biosynthetic process |
| P36542 | ATP5F1C | GO:0006754 | ATP biosynthetic process |
| P48047 | ATP5PO | GO:0006754 | ATP biosynthetic process |
| P56134 | ATP5MF | GO:0006754 | ATP biosynthetic process |
| P56385 | ATP5ME | GO:0006754 | ATP biosynthetic process |
| P04075 | ALDOA | GO:0006754 | ATP biosynthetic process |
| O00483 | NDUFA4 | GO:0045333 | cellular respiration |
| P09669 | COX6C | GO:0045333 | cellular respiration |
| P10606 | COX5B | GO:0045333 | cellular respiration |
| P13073 | COX4I1 | GO:0045333 | cellular respiration |
| P14406 | COX7A2 | GO:0045333 | cellular respiration |
| P14927 | UQCRB | GO:0045333 | cellular respiration |
| P15954 | COX7C | GO:0045333 | cellular respiration |
| P20674 | COX5A | GO:0045333 | cellular respiration |
| P28331 | NDUFS1 | GO:0045333 | cellular respiration |
| P31930 | UQCRC1 | GO:0045333 | cellular respiration |
| P99999 | CYCS | GO:0045333 | cellular respiration |
| P0DP23;P0DP24;P0D | CALM1, CALM2, CAL | GO:0060291 | long-term synaptic potentiation |
| P51608 | MECP2 | GO:0060291 | long-term synaptic potentiation |
| Q13554 | CAMK2B | GO:0060291 | long-term synaptic potentiation |
| Q13555 | CAMK2G | GO:0060291 | long-term synaptic potentiation |
| Q9UQM7 | CAMK2A | GO:0060291 | long-term synaptic potentiation |
| P04350 | TUBB4A | GO:0000226 | microtubule cytoskeleton organization |
| P04406 | GAPDH | GO:0000226 | microtubule cytoskeleton organization |

| | | | |
|---|---|---|---|
| P07437 | TUBB | GO:0000226 | microtubule cytoskeleton organization |
| P0DPH7;P0DPH8 | TUBA3C, TUBA3D | GO:0000226 | microtubule cytoskeleton organization |
| P11137 | MAP2 | GO:0000226 | microtubule cytoskeleton organization |
| P18583 | SON | GO:0000226 | microtubule cytoskeleton organization |
| P46821 | MAP1B | GO:0000226 | microtubule cytoskeleton organization |
| P68363 | TUBA1B | GO:0000226 | microtubule cytoskeleton organization |
| P68366 | TUBA4A | GO:0000226 | microtubule cytoskeleton organization |
| P68371 | TUBB4B | GO:0000226 | microtubule cytoskeleton organization |
| Q00535 | CDK5 | GO:0000226 | microtubule cytoskeleton organization |
| Q13509 | TUBB3 | GO:0000226 | microtubule cytoskeleton organization |
| Q13885 | TUBB2A | GO:0000226 | microtubule cytoskeleton organization |
| Q9BUF5 | TUBB6 | GO:0000226 | microtubule cytoskeleton organization |
| Q9NSV4 | DIAPH3 | GO:0000226 | microtubule cytoskeleton organization |
| Q9NY65 | TUBA8 | GO:0000226 | microtubule cytoskeleton organization |
| Q9NZT1 | CALML5 | GO:0000226 | microtubule cytoskeleton organization |
| P04350 | TUBB4A | GO:0007017 | microtubule-based process |
| P07437 | TUBB | GO:0007017 | microtubule-based process |
| P0DPH7;P0DPH8 | TUBA3C, TUBA3D | GO:0007017 | microtubule-based process |
| P68363 | TUBA1B | GO:0007017 | microtubule-based process |
| P68366 | TUBA4A | GO:0007017 | microtubule-based process |
| P68371 | TUBB4B | GO:0007017 | microtubule-based process |
| Q13509 | TUBB3 | GO:0007017 | microtubule-based process |
| Q13885 | TUBB2A | GO:0007017 | microtubule-based process |
| Q9BUF5 | TUBB6 | GO:0007017 | microtubule-based process |
| Q9NY65 | TUBA8 | GO:0007017 | microtubule-based process |
| O00483 | NDUFA4 | GO:0006123 | mitochondrial electron transport |
| P09669 | COX6C | GO:0006123 | mitochondrial electron transport |
| P10606 | COX5B | GO:0006123 | mitochondrial electron transport |
| P13073 | COX4I1 | GO:0006123 | mitochondrial electron transport |
| P14406 | COX7A2 | GO:0006123 | mitochondrial electron transport |
| P15954 | COX7C | GO:0006123 | mitochondrial electron transport |
| P20674 | COX5A | GO:0006123 | mitochondrial electron transport |
| P99999 | CYCS | GO:0006123 | mitochondrial electron transport |
| P04350 | TUBB4A | GO:0000278 | mitotic cell cycle |
| P07437 | TUBB | GO:0000278 | mitotic cell cycle |
| P0DPH7;P0DPH8 | TUBA3C, TUBA3D | GO:0000278 | mitotic cell cycle |
| P21127 | CDK11B | GO:0000278 | mitotic cell cycle |
| P62826 | RAN | GO:0000278 | mitotic cell cycle |
| P68363 | TUBA1B | GO:0000278 | mitotic cell cycle |
| P68366 | TUBA4A | GO:0000278 | mitotic cell cycle |
| P68371 | TUBB4B | GO:0000278 | mitotic cell cycle |
| Q00610 | CLTC | GO:0000278 | mitotic cell cycle |
| Q13123 | IK | GO:0000278 | mitotic cell cycle |
| Q13509 | TUBB3 | GO:0000278 | mitotic cell cycle |

| | | | |
|---|---|---|---|
| Q13885 | TUBB2A | GO:0000278 | mitotic cell cycle |
| Q9BUF5 | TUBB6 | GO:0000278 | mitotic cell cycle |
| Q9NY65 | TUBA8 | GO:0000278 | mitotic cell cycle |
| Q9UQE7 | SMC3 | GO:0000278 | mitotic cell cycle |
| P62714;P67775 | PPP2CB, PPP2CA | GO:0000278 | mitotic cell cycle |
| P09669 | COX6C | GO:0006119 | oxidative phosphorylation |
| P10606 | COX5B | GO:0006119 | oxidative phosphorylation |
| P13073 | COX4I1 | GO:0006119 | oxidative phosphorylation |
| P14406 | COX7A2 | GO:0006119 | oxidative phosphorylation |
| P14927 | UQCRB | GO:0006119 | oxidative phosphorylation |
| P15954 | COX7C | GO:0006119 | oxidative phosphorylation |
| P20674 | COX5A | GO:0006119 | oxidative phosphorylation |
| P31930 | UQCRC1 | GO:0006119 | oxidative phosphorylation |
| P36542 | ATP5F1C | GO:0006119 | oxidative phosphorylation |
| P43246 | MSH2 | GO:0006119 | oxidative phosphorylation |
| O75947 | ATP5PD | GO:0015986 | proton motive force-driven ATP synthesis |
| P06576 | ATP5F1B | GO:0015986 | proton motive force-driven ATP synthesis |
| P24539 | ATP5PB | GO:0015986 | proton motive force-driven ATP synthesis |
| P25705 | ATP5F1A | GO:0015986 | proton motive force-driven ATP synthesis |
| P30049 | ATP5F1D | GO:0015986 | proton motive force-driven ATP synthesis |
| P36542 | ATP5F1C | GO:0015986 | proton motive force-driven ATP synthesis |
| P48047 | ATP5PO | GO:0015986 | proton motive force-driven ATP synthesis |
| P56134 | ATP5MF | GO:0015986 | proton motive force-driven ATP synthesis |
| P56385 | ATP5ME | GO:0015986 | proton motive force-driven ATP synthesis |
| Q96IX5 | ATP5MK | GO:0015986 | proton motive force-driven ATP synthesis |
| O75494 | SRSF10 | GO:0048024 | regulation of mRNA splicing, via spliceosome |
| O75525 | KHDRBS3 | GO:0048024 | regulation of mRNA splicing, via spliceosome |
| P18583 | SON | GO:0048024 | regulation of mRNA splicing, via spliceosome |
| P61978 | HNRNPK | GO:0048024 | regulation of mRNA splicing, via spliceosome |
| P84103 | SRSF3 | GO:0048024 | regulation of mRNA splicing, via spliceosome |
| Q07666 | KHDRBS1 | GO:0048024 | regulation of mRNA splicing, via spliceosome |
| Q14498 | RBM39 | GO:0048024 | regulation of mRNA splicing, via spliceosome |
| Q96MU7 | YTHDC1 | GO:0048024 | regulation of mRNA splicing, via spliceosome |
