## Supplementary material for "Single-cell nanodroplet processing proteomics pipeline for analysis of human-derived microglia": Supp_3.pdf

| Accession | Gene | GO_ID | GO_Term |
|---|---|---|---|
| P00751 | CFB | GO:0006956 | complement activation |
| P01024 | C3 | GO:0006956 | complement activation |
| P01031 | C5 | GO:0006956 | complement activation |
| P02745 | C1QA | GO:0006956 | complement activation |
| P02748 | C9 | GO:0006956 | complement activation |
| P05156 | CFI | GO:0006956 | complement activation |
| P08603 | CFH | GO:0006956 | complement activation |
| P09871 | C1S | GO:0006956 | complement activation |
| P0C0L4 | C4A | GO:0006956 | complement activation |
| P0C0L5 | C4B | GO:0006956 | complement activation |
| P10909 | CLU | GO:0006956 | complement activation |
| P13987 | CD59 | GO:0006956 | complement activation |
| P00751 | CFB | GO:0006957 | complement activation, alternative pathway |
| P01024 | C3 | GO:0006957 | complement activation, alternative pathway |
| P01031 | C5 | GO:0006957 | complement activation, alternative pathway |
| P02748 | C9 | GO:0006957 | complement activation, alternative pathway |
| P08603 | CFH | GO:0006957 | complement activation, alternative pathway |
| P01024 | C3 | GO:0006958 | complement activation, classical pathway |
| P01031 | C5 | GO:0006958 | complement activation, classical pathway |
| P01857 | IGHG1 | GO:0006958 | complement activation, classical pathway |
| P01859 | IGHG2 | GO:0006958 | complement activation, classical pathway |
| P01861 | IGHG4 | GO:0006958 | complement activation, classical pathway |
| P01876 | IGHA1 | GO:0006958 | complement activation, classical pathway |
| P01877 | IGHA2 | GO:0006958 | complement activation, classical pathway |
| P02745 | C1QA | GO:0006958 | complement activation, classical pathway |
| P02747 | C1QC | GO:0006958 | complement activation, classical pathway |
| P02748 | C9 | GO:0006958 | complement activation, classical pathway |

| | | | |
|---|---|---|---|
| P04003 | C4BPA | GO:0006958 | complement activation, classical pathway |
| P05156 | CFI | GO:0006958 | complement activation, classical pathway |
| P09871 | C1S | GO:0006958 | complement activation, classical pathway |
| P0C0L4 | C4A | GO:0006958 | complement activation, classical pathway |
| P0C0L5 | C4B | GO:0006958 | complement activation, classical pathway |
| P10909 | CLU | GO:0006958 | complement activation, classical pathway |
| Q07021 | C1QBP | GO:0006958 | complement activation, classical pathway |
| Q16181 | SEPTIN7 | GO:0061640 | cytoskeleton-dependent cytokinesis |
| Q9NVA2 | SEPTIN11 | GO:0061640 | cytoskeleton-dependent cytokinesis |
| Q9UHD8 | SEPTIN9 | GO:0061640 | cytoskeleton-dependent cytokinesis |
| O75935 | DCTN3 | GO:0061640 | cytoskeleton-dependent cytokinesis |
| P53990 | IST1 | GO:0061640 | cytoskeleton-dependent cytokinesis |
| Q15019 | SEPTIN2 | GO:0061640 | cytoskeleton-dependent cytokinesis |
| B5ME19;Q99613 | EIF3CL, EIF3C | GO:0001732 | formation of cytoplasmic translation initiation complex |
| O00303 | EIF3F | GO:0001732 | formation of cytoplasmic translation initiation complex |
| O15371 | EIF3D | GO:0001732 | formation of cytoplasmic translation initiation complex |
| O75821 | EIF3G | GO:0001732 | formation of cytoplasmic translation initiation complex |
| O75822 | EIF3J | GO:0001732 | formation of cytoplasmic translation initiation complex |
| P55010 | EIF5 | GO:0001732 | formation of cytoplasmic translation initiation complex |

| | | | |
|---|---|---|---|
| P55884 | EIF3B | GO:0001732 | formation of cytoplasmic translation initiation complex |
| P60228 | EIF3E | GO:0001732 | formation of cytoplasmic translation initiation complex |
| Q14152 | EIF3A | GO:0001732 | formation of cytoplasmic translation initiation complex |
| Q9UBQ5 | EIF3K | GO:0001732 | formation of cytoplasmic translation initiation complex |
| Q9Y262 | EIF3L | GO:0001732 | formation of cytoplasmic translation initiation complex |
| O15372 | EIF3H | GO:0001732 | formation of cytoplasmic translation initiation complex |
| Q7L2H7 | EIF3M | GO:0001732 | formation of cytoplasmic translation initiation complex |
| O95445 | APOM | GO:0034384 | high-density lipoprotein particle clearance |
| P02647 | APOA1 | GO:0034384 | high-density lipoprotein particle clearance |
| P02649 | APOE | GO:0034384 | high-density lipoprotein particle clearance |
| P02655 | APOC2 | GO:0034384 | high-density lipoprotein particle clearance |
| P02652 | APOA2 | GO:0034384 | high-density lipoprotein particle clearance |
| O95445 | APOM | GO:0034375 | high-density lipoprotein particle remodeling |
| P02647 | APOA1 | GO:0034375 | high-density lipoprotein particle remodeling |
| P02649 | APOE | GO:0034375 | high-density lipoprotein particle remodeling |
| P02654 | APOC1 | GO:0034375 | high-density lipoprotein particle remodeling |
| P02656 | APOC3 | GO:0034375 | high-density lipoprotein particle remodeling |
| P06727 | APOA4 | GO:0034375 | high-density lipoprotein particle remodeling |

| | | | |
|---|---|---|---|
| P02652 | APOA2 | GO:0034375 | high-density lipoprotein particle remodeling |
| P10606 | COX5B | GO:0006123 | mitochondrial electron transport, cytochrome c to oxygen |
| P99999 | CYCS | GO:0006123 | mitochondrial electron transport, cytochrome c to oxygen |
| P00403 | MT-CO2 | GO:0006123 | mitochondrial electron transport, cytochrome c to oxygen |
| P09669 | COX6C | GO:0006123 | mitochondrial electron transport, cytochrome c to oxygen |
| P13073 | COX4I1 | GO:0006123 | mitochondrial electron transport, cytochrome c to oxygen |
| O00483 | NDUFA4 | GO:0006123 | mitochondrial electron transport, cytochrome c to oxygen |
| P14406 | COX7A2 | GO:0006123 | mitochondrial electron transport, cytochrome c to oxygen |
| P00734 | F2 | GO:0051918 | negative regulation of fibrinolysis |
| P00747 | PLG | GO:0051918 | negative regulation of fibrinolysis |
| P02749 | APOH | GO:0051918 | negative regulation of fibrinolysis |
| P04004 | VTN | GO:0051918 | negative regulation of fibrinolysis |
| P04196 | HRG | GO:0051918 | negative regulation of fibrinolysis |
| P08697 | SERPINF2 | GO:0051918 | negative regulation of fibrinolysis |
| P05121 | SERPINE1 | GO:0051918 | negative regulation of fibrinolysis |
| O60220 | TIMM8A | GO:0045039 | protein insertion into mitochondrial inner membrane |
| O94826 | TOMM70 | GO:0045039 | protein insertion into mitochondrial inner membrane |

| | | | |
|---|---|---|---|
| Q53H12 | AGK | GO:0045039 | protein insertion into mitochondrial inner membrane |
| Q9Y5J7 | TIMM9 | GO:0045039 | protein insertion into mitochondrial inner membrane |
| Q9Y5J9 | TIMM8B | GO:0045039 | protein insertion into mitochondrial inner membrane |
| Q9Y5L4 | TIMM13 | GO:0045039 | protein insertion into mitochondrial inner membrane |
| O43809 | NUDT21 | GO:0051262 | protein tetramerization |
| P34897 | SHMT2 | GO:0051262 | protein tetramerization |
| P04637 | TP53 | GO:0051262 | protein tetramerization |
| Q15070 | OXA1L | GO:0051262 | protein tetramerization |
| Q16630 | CPSF6 | GO:0051262 | protein tetramerization |
| Q8N684 | CPSF7 | GO:0051262 | protein tetramerization |
| O15371 | EIF3D | GO:0075525 | viral translational termination-reinitiation |
| O75821 | EIF3G | GO:0075525 | viral translational termination-reinitiation |
| P55884 | EIF3B | GO:0075525 | viral translational termination-reinitiation |
| Q14152 | EIF3A | GO:0075525 | viral translational termination-reinitiation |
| Q9Y262 | EIF3L | GO:0075525 | viral translational termination-reinitiation |
